# The large loop-8 insert in *Saccharomyces cerevisiae* kinesin-5 Cin8 is necessary and sufficient to promote noncanonical microtubule interactions

**DOI:** 10.1101/525329

**Authors:** Kayla M. Bell, Jared C. Cochran

## Abstract

*Saccharomyces cerevisiae* kinesin-5 Cin8 displays unconventional biochemical behavior including bidirectional motility and ability to bind multiple motor domains per αβ tubulin dimer in the microtubule lattice. Previous research suggested that a large loop-8 insert near the microtubule binding interface of Cin8 was critical for its noncanonical microtubule binding behavior. Here we utilized mutagenesis, thermodynamic, and kinetic assays to further understand the mechanism for how this loop-8 insert promotes super-stoichiometric microtubule binding in Cin8. This loop-8 insert that interrupts the conserved β5a/b hairpin was swapped between Cin8, Eg5 (KIF11, a human kinesin-5) and Kip1 (another *S. cerevisiae* kinesin-5). Cin8 with the loop-8 insert from Eg5 (Cin8-EL8) binds one motor per tubulin dimer, whereas Eg5 with the loop-8 insert from Cin8 (Eg5-CL8) binds approximately 2-4 motors per tubulin dimer. Eg5-CL8 bound the canonical and noncanonical sites on the microtubule lattice with weakened oligomerization between motors, while Cin8-EL8 showed only canonical site binding. These results demonstrate that the large loop-8 insert in Cin8 is necessary and sufficient to promote noncanonical microtubule binding behavior.

## INTRODUCTION

In budding yeast (*Saccharomyces cerevisiae*), Cin8 and Kip1 belong to the kinesin-5 subfamily and share overlapping roles in microtubule cross-linking, antiparallel microtubule sliding, and regulating microtubule dynamics for spindle assembly and maintenance (1–8). Full length Cin8 and Kip1 have been shown to move in both directions along the microtubule, both *in vitro* and *in vivo*, and switches directionality based on ionic strength, relative microtubule orientation (i.e. parallel *versus* antiparallel interaction), and motor density (9–12). Additionally, bidirectional movement in a stable dimeric Cin8/kinesin-1 chimera (consisting of the Cin8 catalytic core and neck linker fused to the kinesin-1 stalk) has supported the hypothesis that Cin8 has intrinsic bidirectionality within the N-terminal motor domain, with directional switching regulated by the C-terminal tail domain (13). The mechanism of how yeast kinesin-5 motors achieve this bidirectional motility is still not well understood.

Characterization of the microtubule binding and ATPase activity of monomeric Cin8 revealed a novel interaction with the microtubule such that 4 motor domains bound per tubulin dimer in the microtubule lattice (14). We proposed that loop-8 played a critical role in 1) binding at the canonical and (yet structurally uncharacterized) noncanonical microtubule sites and 2) oligomerization of the Cin8 motors at these microtubule sites (14). Loop-8 is part of the kinesin microtubule binding interface, and mutations in this loop can modulate microtubule binding interaction (15–20). Furthermore, loop-8 has been shown to influence directional switching in Cin8 such that when the large (99-residue) insert in loop-8 was replaced with the homologous (6-residue) insert from Kip1, the Cin8 motor lost its ability to bind the microtubule at higher ionic strength, and the switch from minus-stepping at high ionic strength to plus-stepping at low ionic strength occurred at a much lower salt concentration (10,14).

In this study, we utilized site-directed mutagenesis, steady-state kinetics, and equilibrium cosedimentation assays to dissect the mechanism for how the large loop-8 insert modulates microtubule binding of Cin8. To this end, we cloned, expressed and purified chimeric motors of Cin8, Eg5 and Kip1 in which their loop-8 inserts were swapped, creating a monomeric Eg5 motor with the loop-8 insert of Cin8 (Eg5-CL8), a monomeric Cin8 motor with the loop-8 insert of Eg5 (Cin8-EL8), and a monomeric Eg5 motor domain with the loop-8 insert for Kip1 (Eg5-KL8). A monomeric Cin8 motor with the loop-8 insert of Kip1 (Cin8-KL8, also known as Cin8-Δ99) was previously characterized (10,14). Cin8-EL8 and Eg5-KL8 showed canonical binding, while Eg5-CL8 showed both canonical and noncanonical binding, but with weakened oligomerization on these microtubule sites.

## RESULTS

### Rational design of Cin8 and Eg5 mutant motors

We cloned, expressed and purified new chimeric constructs of Cin8, Eg5, and Kip1 motor domains (Fig. 1a): Cin8 containing the loop-8 insert from Eg5 (Cin8-EL8 and Cin8-EL8-trx), Eg5 containing the loop-8 insert from Cin8 (Eg5-CL8 and Eg5-CL8-MBP), and Eg5 containing the loop-8 insert from Kip1 (Eg5-KL8). Based on a Clustal Omega alignment of kinesin-5 motors (Fig. 1b) and a previously published Cin8-KL8 mutant (10), we swapped the loop-8 regions for the three motors that aligned with the large loop-8 insert from Cin8 (from residues 261 to 359). These correspond to resides 187 to 193 in monomeric Eg5 (21), and 186 to 284 in monomeric Cin8 (14).

**Figure 1.**
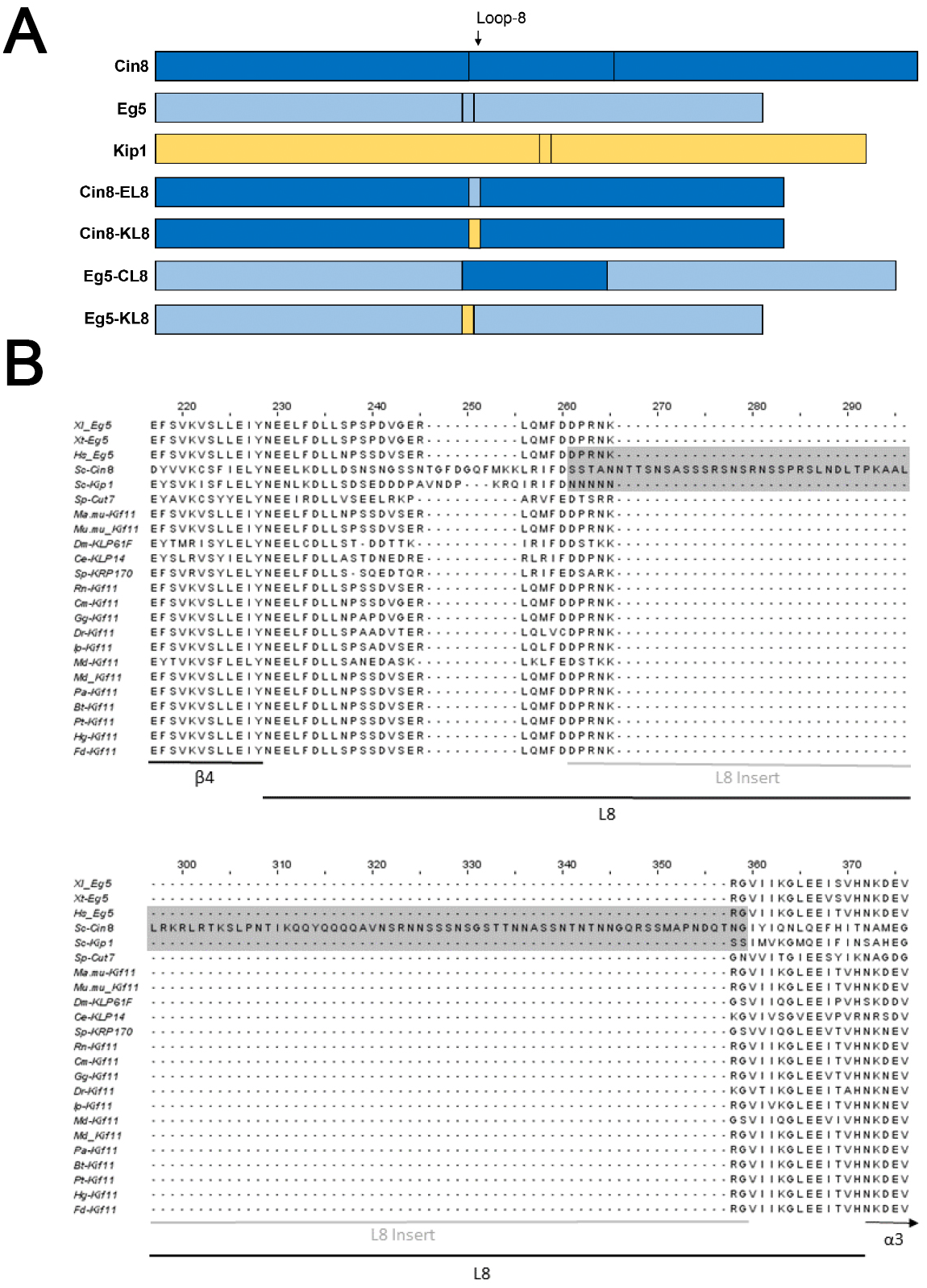
Construct designs and alignment of kinesin-5 loop-8 sequences. (**A**) Depiction of the design of the motor constructs used, showing the wild type constructs and the swapped loop-8 regions (not to scale). Regions from Cin8 are shown in purple, regions from Eg5 are shown in red, and regions from Kip1 are shown in light blue. Yellow marks the approximate location of the SCRK mutations in Eg5-SCRK. (**B**) ClustalOmega alignment of kinesin-5 motors showing the loop-8 region and flanking sequences. The three bidirectional motors, Cin8, Kip1 and Cut7, show a large amount of variation in their loop-8 regions and the sequences immediately flanking it. The loop-8 insert swapped in the mutant motor proteins in highlighted in grey.

### Swapped loop-8 motors have typical ATPase activity like Eg5 and Cin8

We performed a thorough steady state kinetic analysis on mutant motors, both tagless and tagged constructs (Fig. 2; Table 1), and compared them with the wild type Eg5 and Cin8 motor domains (Eg5-WT and Cin8-WT) to see if the loop-8 insert influenced microtubule binding affinity, ATP binding affinity, and ATPase activity. All three constructs had typical microtubule and ATP binding and ATPase rates for a kinesin-5 motor.

**TABLE 1.**
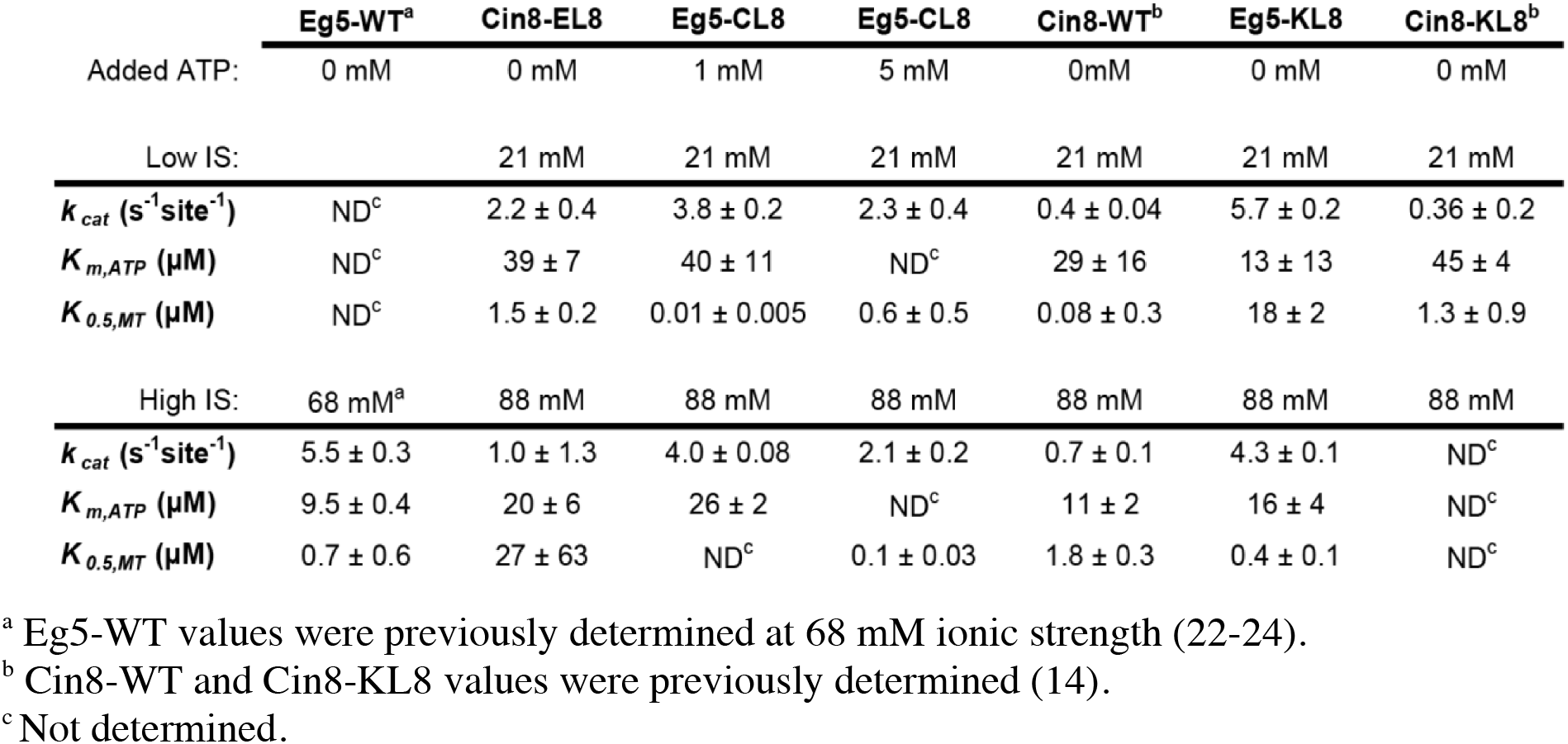
Steady-state kinetic constants for kinesin-5 motors.

|  | Eg5-WT<sup>a</sup> | Cin8-EL8 | Eg5-CL8 | Eg5-CL8 | Cin8-WT<sup>b</sup> | Eg5-KL8 | Cin8-KL8<sup>b</sup> |
| --- | --- | --- | --- | --- | --- | --- | --- |
| Added ATP: | 0 mM | 0 mM | 1 mM | 5 mM | 0mM | 0 mM | 0 mM |
| Low IS: |  | 21 mM | 21 mM | 21 mM | 21 mM | 21 mM | 21 mM |
| $k_{cat} (\text{s}^{-1} \text{ site}^{-1})$ | ND<sup>c</sup> | $2.2 \pm 0.4$ | $3.8 \pm 0.2$ | $2.3 \pm 0.4$ | $0.4 \pm 0.04$ | $5.7 \pm 0.2$ | $0.36 \pm 0.2$ |
| $K_{m,ATP} (\mu\text{M})$ | ND<sup>c</sup> | $39 \pm 7$ | $40 \pm 11$ | ND<sup>c</sup> | $29 \pm 16$ | $13 \pm 13$ | $45 \pm 4$ |
| $K_{0.5,MT} (\mu\text{M})$ | ND<sup>c</sup> | $1.5 \pm 0.2$ | $0.01 \pm 0.005$ | $0.6 \pm 0.5$ | $0.08 \pm 0.3$ | $18 \pm 2$ | $1.3 \pm 0.9$ |
| High IS: | 68 mM<sup>a</sup> | 88 mM | 88 mM | 88 mM | 88 mM | 88 mM | 88 mM |
| $k_{cat} (\text{s}^{-1} \text{ site}^{-1})$ | $5.5 \pm 0.3$ | $1.0 \pm 1.3$ | $4.0 \pm 0.08$ | $2.1 \pm 0.2$ | $0.7 \pm 0.1$ | $4.3 \pm 0.1$ | ND<sup>c</sup> |
| $K_{m,ATP} (\mu\text{M})$ | $9.5 \pm 0.4$ | $20 \pm 6$ | $26 \pm 2$ | ND<sup>c</sup> | $11 \pm 2$ | $16 \pm 4$ | ND<sup>c</sup> |
| $K_{0.5,MT} (\mu\text{M})$ | $0.7 \pm 0.6$ | $27 \pm 63$ | ND<sup>c</sup> | $0.1 \pm 0.03$ | $1.8 \pm 0.3$ | $0.4 \pm 0.1$ | ND<sup>c</sup> |

<sup>a</sup> Eg5-WT values were previously determined at 68 mM ionic strength (22-24).
<sup>b</sup> Cin8-WT and Cin8-KL8 values were previously determined (14).
<sup>c</sup> Not determined.

**Figure 2.**
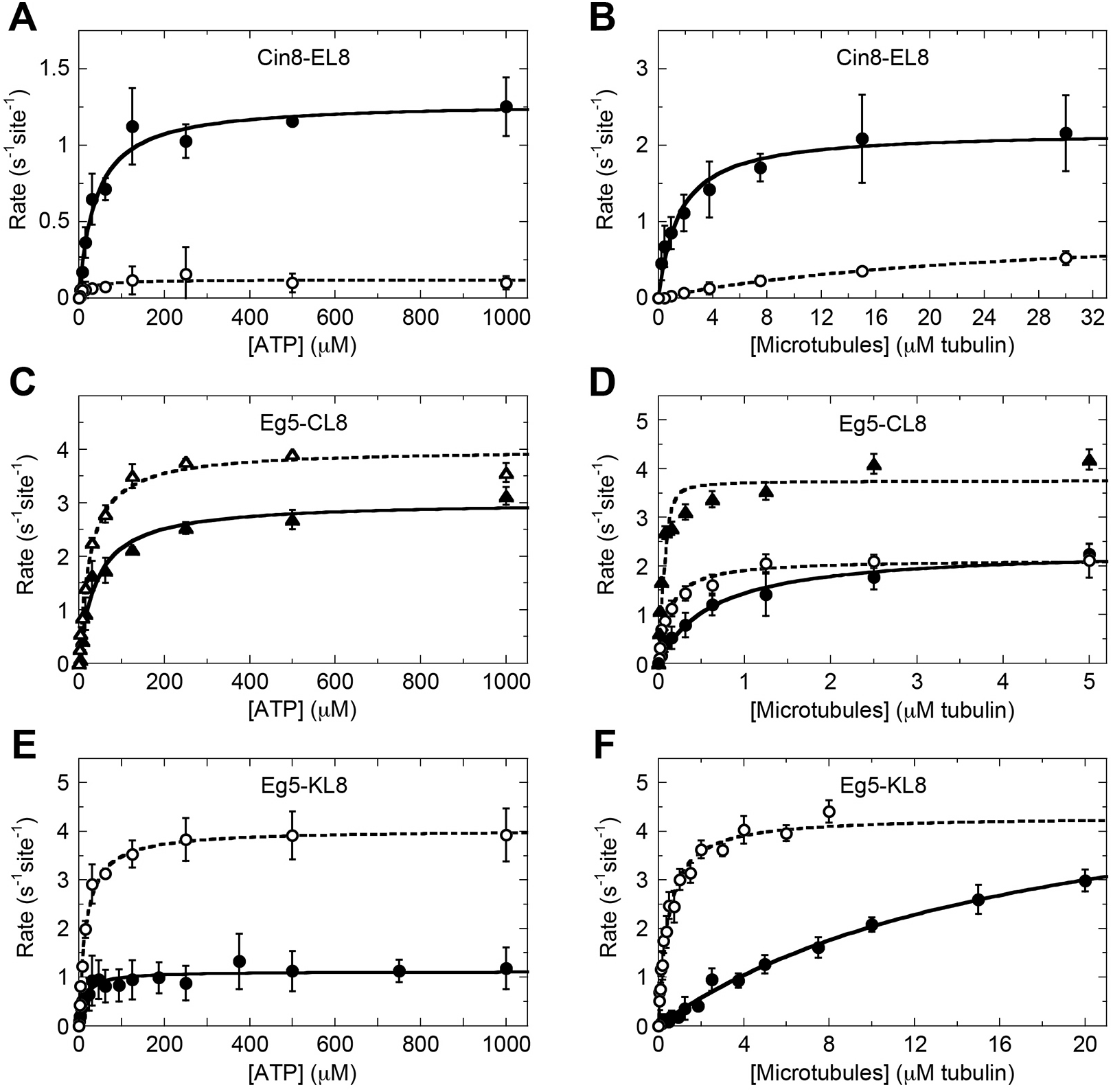
Steady-state kinetics of mutant kinesins indicate typical kinesin-5 ATPase activity. (**A**) ATP-stimulated activity for Cin8-EL8 at low (closed circles) and high (open circles) ionic strength. Final concentrations are 0.1 μM Cin8-EL8 (tagless), 0-1000 μM ATP and 5 μM MT. Low IS, *K_m, ATP_* = 39 ± 7 μM. High IS, *K_m, ATP_* = 20 ± 9 μM. (**B**) Microtubule-stimulated ATPase activity for Cin8-EL8 at low (closed circles) and high (open circles) ionic strength. Final concentrations are 0.1 μM Cin8-EL8 (tagless), 0-30 μM MTs, 1 mM ATP. Low IS, *K_cat_* = 2.2 ±0.1 s^-2^site^-2^ and *k_0.5,MT_* = 1.5 ±0.3 μM. High IS, *k_cat_* = 1.0 ±0.07 s^-1^site^-1^ and *K_0.5,MT_* = 27 ± 3.1 μM. (**C**) ATP-stimulated activity for Eg5-CL8 at low (closed triangles) and high (open triangles) ionic strength. Final concentrations are 0.05 μM Eg5-CL8 (tagless), 0-1000 μM ATP and 2 μM MT. Low IS, *k_m,ATP_* = 40 ± 7 μM. High IS, *K_m, ATP_* = 26 ± 3 μM and *k_cat_* = 4.0 ±0.1 s^-1^site^-1^. (**D**) Microtubule-stimulated ATPase activity for Eg5-CL8 at low (closed circles) and high (open circles) ionic strength with 5 niM ATP, and at low ionic strength with 1 niM ATP (closed triangles). Final concentrations at 5 mM ATP are 0.025 μM Eg5-CL8 (tagless), 0-5 μM MTs, 5 niM ATP. Final concentrations at 1 mM ATP are 0.1 μM Eg5-CL8 (tagless), 0-5 μM MTs, 1 mM ATP. Low IS, 5 mM ATP, *k_cat_* = 2.3 ± 0.1 s^-1^site^-1^ and *K_0.5, MT_* = 0.6 ± 0.1 μM. High IS, 5 mM ATP, *k_cat_* = 2.1 ± 0.07 s^-1^site^-1^ and *K_0.5, MT_* = 0.1 ± 0.02 μM. Low IS, 1 mM ATP, *k_cat_* = 3.8 ± 0.2 s^-1^site^-1^ and *K_0.5, MT_* = 0.01 ± 0.01 μM. (**E**) ATP-stimulated activity for Eg5-KL8 at low (closed circles) and high (open circles) ionic strength. Final concentrations are 0.05 μM Eg5-KL8, 0-1000 μM ATP and 10 μM MT. Low IS, *K_m, ATP_* = 13 ± 2 μM. High IS, *K_m,ATP_* = 16 ± 1 μM. (**F**) Microtubule-stimulated ATPase activity for Eg5-KL8 at low (closed circles) and high (open circles) ionic strength. Final concentrations are 0.05 μM Eg5-KL8, 0-20 μM MTs (Low IS), 0-8 μM MTs (High IS) 1 mM ATP. Low IS, *k_cat_* = 5.7 ± 0.5 s^-1^site^-1^ and *K_0.5, MT_* = 18 ± 2.4 μM. High IS, *k_cat_* = 4.3 ± 0.1 s^-1^site^-1^ and *K_0.5, MT_* = 0.4 ± 0.04 μM.

Cin8-EL8 showed relatively tight ATP binding that was comparable at low (*K_m,ATP_* = 39 ± 7 μM) and high (*K_m,ATP_* = 20 ± 6 μM) ionic strength (Fig. 2a). It showed tighter microtubule binding at low ionic strength (*K_0.5,MT_* = 1.5 ± 0.2 μM), and approximately 11-fold weaker binding at high ionic strength (*K_0.5, MT_* = 27 ± 63 μM) (Fig. 2b). Cin8-EL8 had a *k_cat_* slower than Eg5-WT but faster than Cin8-WT. It also showed an approximately 2-fold difference between its *k_cat_* at low (*k_cat_* = 2.2 ± 0.4 s^-1^ site^-1^) and high (*k_cat_* = 1.0 ± 1.3 s^-1^ site^-1^) ionic strength.

Under nanomolar motor and 1 mM ATP conditions, Eg5-CL8 had robust ATPase activity with a relatively tight ATP binding that was comparable at low (*K_m,ATP_* = 40 ± 11 μM) and high (*K_m,ATP_* = 26 ± 2 μM) ionic strength (Fig 2c), and showed tight binding to microtubules at low ionic strength (*K_m,MT_* = 0.01 ± 0.005 μM) (Fig 2d, Table 1). However, when TEV cleaving the concentrated Eg5-CL8-MBP (40 μM) to get a tagless version for use in steady-state experiments, the Eg5-CL8 (tagless) was observed to precipitate out of the solution, and the visible aggregates would pellet when centrifuged. Addition of 5 mM ATP returned the aggregated Eg5-CL8 (tagless) to solution (data not shown), so steady state experiments for Eg5-CL8 (tagless) were also done in the presence of 5 mM ATP. Eg5-CL8 under these conditions showed tight microtubule binding at both low ionic strength (*K_0.5,MT_* = 0.6 ± 0.5 μM) and high (*K_0.5,MT_* = 0.1 ± 0.03 μM) ionic strength (Fig. 2d). No ATP-dependent data were gathered under these conditions, because in ATP dependent experiments the ATP concentration needs to be varied to determine binding affinity. At 5 mM ATP, the *k_cat_* for Eg5-CL8 was comparable at both low (*k_cat_* = 2.3 ± 0.4 s^-1^ site^-1^) and high (*k_cat_* = 2.1 ± 0.2 s^-1^ site^-1^) ionic strength, and was slower than that of wild type Eg5 (*k_cat_* = 5.5 ± 0.3 s^-1^ site^-1^), but was like the rate for Cin8-EL8. Additionally, this *k_cat_* was approximately half the rate seen for the same Eg5-CL8 construct under 1 mM ATP conditions (Low IS, *k_cat_* = 3.8 ± 0.2 s^-1^ site^-1^. High IS, *k_cat_* = 4.0 ± 0.08 s^-1^ site^-1^) (Fig. 2c).

Eg5-KL8 showed relatively tight ATP binding that was comparable at low (*K_m,ATP_* = 13 ± 13 μM) and high (*K_m,ATP_* = 16 ± 4 μM) ionic strength (Fig. 2e). Surprisingly, it showed tight microtubule binding at high ionic strength (*K_0.5,MT_* = 0.4 ± 0.1 μM), and approximately 45fold weaker binding at low ionic strength (*K_0.5,MT_* = 18 ± 2 μM) (Fig. 2f). Eg5-KL8 had a *k_cat_* like that of Eg5-WT (Eg5-WT, *k_cat_* = 5.5 ± 0.3 s^-1^ site^-1^) and appeared to have a slightly faster *k_cat_* at low ionic strength (k_cat_ = 5.7 ± 0.2 s^-1^ site^-1^) when compared to high ionic strength (*k_cat_* = 4.3 ± 0.06 s^-1^ site^-1^) (Table 1).

### Cin8-EL8 displays stoichiometric interaction with the microtubule like Eg5-WT

The equilibrium binding affinities (*K_d,app_*) for microtubules and the stoichiometry coefficient (*s*) of Cin8-EL8 binding to the tubulin dimer in the microtubule lattice were measured using cosedimentation assays. Cin8-EL8 partitioned with the microtubules with maximal binding occurring at a tubulin to kinesin molar ratio of approximately 1 under low ionic strength conditions (Low IS, *s* = 1.4 ± 0.3. Fig. 3a). At low ionic strength, Cin8-EL8 showed tight microtubule binding, (*K_d, app_* = 0.1 ± 0.04 μM), which weakened 30-fold at high ionic strength (*K_d,app_* = 3.0 ± 0.8 μM) (Table 2). This weakened microtubule binding was significantly more than seen in Cin8-WT or Eg5-WT at high ionic strength.

**TABLE 2.** Microtubule equilibrium binding constants for kinesin-5 motors.

|  | Eg5-WT<sup>b</sup> | Cin8-EL8 | Eg5-CL8 | Cin8-WT | Cin8-WT<sup>b</sup> | Eg5-KL8 |
| --- | --- | --- | --- | --- | --- | --- |
| Added ATP: | 0 mM | 0 mM | 5 mM | 5 mM | 0 mM | 0 mM |
| Low IS: | 21 mM | 21 mM | 21 mM | 21 mM | 21 mM | 21 mM |
| $K_{d,app}$<sup>a</sup> ($\mu\text{M}$) | $0.01 \pm 0.007$ | $0.1 \pm 0.04$ | $0.8 \pm 0.5$ | $0.007 \pm 0.004$ | $0.001 \pm 0.003$ | $0.44 \pm 0.06$ |
| $s$ ($\mu\text{M}$) | $1.1 \pm 0.7$ | $1.4 \pm 0.3$ | $3.3 \pm 0.9$ | $2.2 \pm 0.1$ | $5.1 \pm 4.2$ | ND<sup>c</sup> |
| High IS: | 88 mM | 88 mM | 88 mM |  | 88 mM | 88 mM |
| $K_{d,app}$<sup>a</sup> ($\mu\text{M}$) | $0.5 \pm 0.2$ | $2.9 \pm 0.2$ | $0.04 \pm 0.004$ | ND<sup>c</sup> | $0.1 \pm 0.4$ | $0.1 \pm 0.06$ |
| $s$ ($\mu\text{M}$) | $1.4 \pm 0.7$ | ND<sup>c</sup> | $1.2 \pm 0.2$ | ND<sup>c</sup> | $4.0 \pm 3.3$ | $1.3 \pm 0.4$ |

<sup>a</sup> Average fit constants when stoichiometric coefficient ( $s$ ) held fixed at $s=1$ for Eg5-WT, Cin8-EL8, and Eg5-KL8, and floated for Cin8-WT (Cin8-trx) and Eg5-CL8.
<sup>b</sup> $K_{d,app}$ and $s$ values were determined previously for Eg5 and Cin8-trx (14).
<sup>c</sup> Not determined.

**Figure 3.**
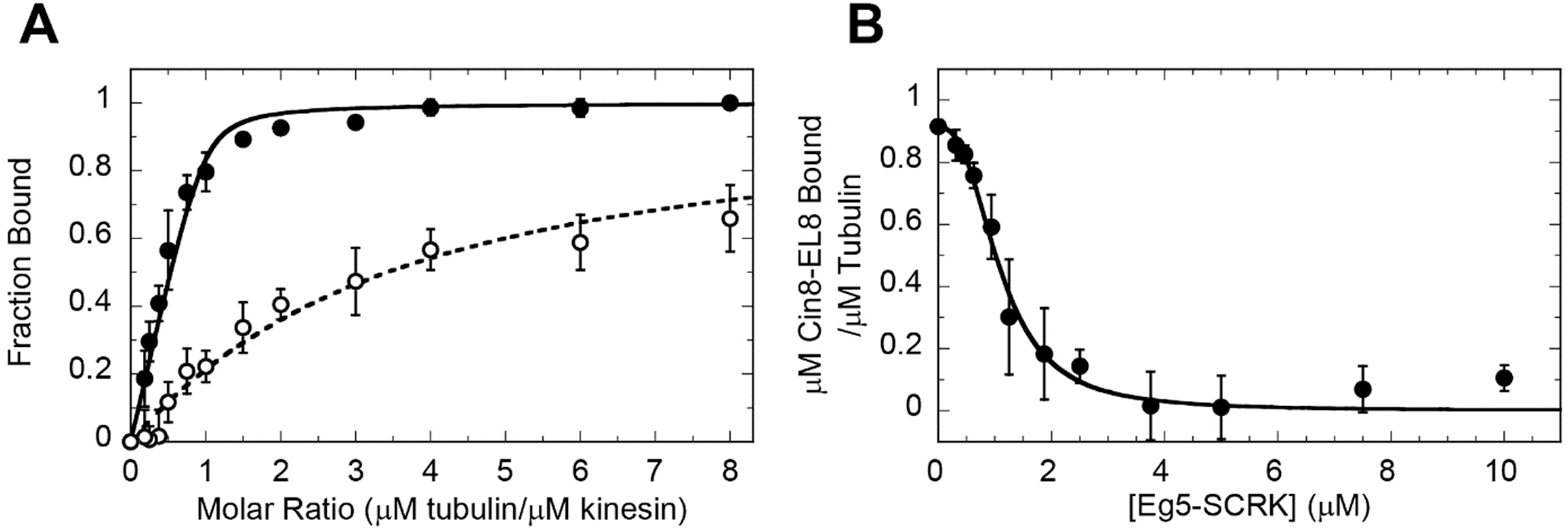
Cosedimentation and competition assays of Cin8-EL8 reveal only stoichiometric and canonical binding to the microtubules. (**A**) Cosedimentation of Cin8-EL8 (tagless) at low (closed circles) and high (open circles) ionic strength. Final concentrations are 2 μM Cin8-EL8 (tagless), 0-8 μM MTs. Low IS, *K_d,app_* = 0.1 ± 0.03 μM and s = 1.4 ± 0.08. High IS, *K_d,app_* = 2.9 ± 0.2 μM. (**B**) Plot of average fraction Cin8-EL8 (tagless) bound to the microtubule at increasing Eg5-SCRK concentrations. Final concentrations are 1 μM Cin8-EL8 (tagless), 1 μM MTs, 0-15 μMEg5-SCRK.

To confirm whether or not the *s*-value of approximately 1 for Cin8-EL8 actually indicates stoichiometric binding, or was just a product of the weakened microtubule binding, a competition assay between Cin8-EL8 and Eg5-SCRK (and mutant deficient for ATP hydrolysis; (25)) was carried out using the NADH coupled assay to quantify the fraction of Cin8-EL8 bound to the microtubule at increasing concentrations of Eg5-SCRK (Fig. 3b). The Eg5-SCRK was observed to completely displace Cin8-EL8 from the microtubule at a 4:1 ratio of Eg5-SCRK to Cin8-EL8. This strongly indicated that Cin8-EL8 does not exhibit the super-stoichiometric binding seen in Cin8-WT. Additionally, under increasing Cin8-EL8 to microtubule ratios above a stoichiometric ratio, cosedimentation of the Cin8-EL8 with the microtubule showed a hyperbolic saturation of Cin8-EL8 bound to the microtubule at approximately 1 motor bound per tubulin dimer (Fig. 4).

**Figure 4.**
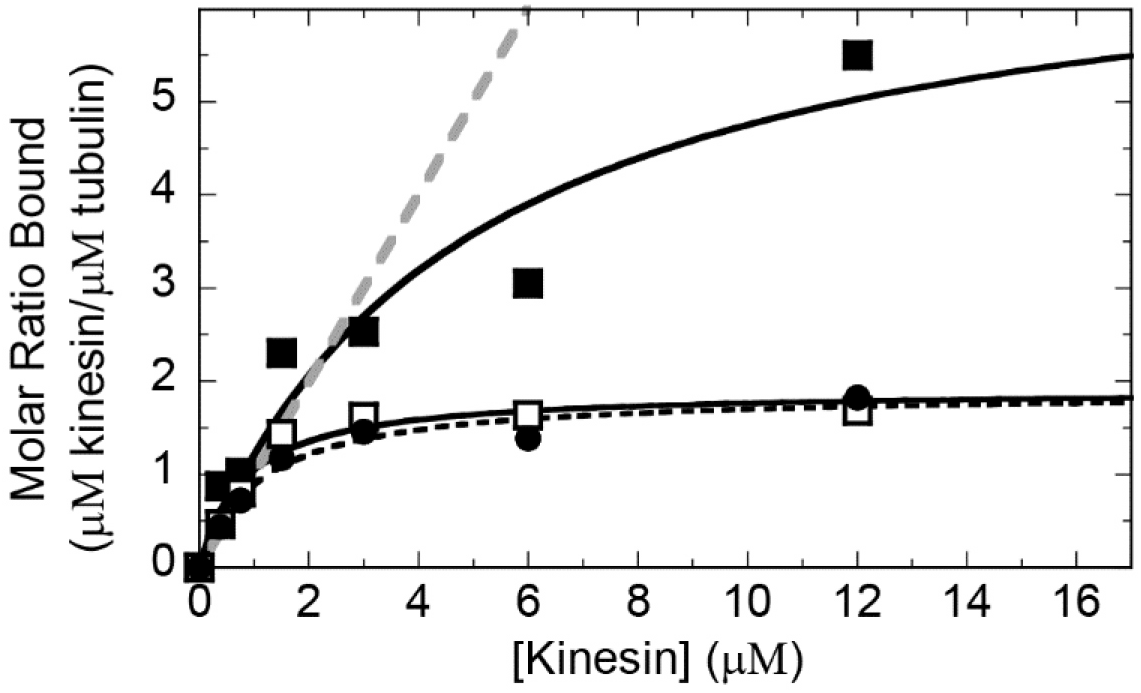
Increasing concentrations of kinesin to a fixed concentration of microtubules shows a saturation of kinesin motors binding to the microtubule. Cosedimentation of Cin8-EL8 (closed circle, dashed line), Eg5-CL8 (closed square, solid line), Eg5-CL8-MBP (open squares, solid line) at increasing super-stoichiometric ratios of kinesin to a fixed amount of microtubules at low ionic strength. Data were fit to a hyperbola. Grey dashed line represents a linear model based on the BET adsorption theory. Final concentrations: 0-12 μM kinesin, 1 μM MT.

### Eg5-CL8 shows super-stoichiometric interaction with the microtubule like Cin8-WT

The *K_d,app_* and s-value were also determined for Eg5-CL8 and Eg5-CL8-MBP by cosedimentation assays (Fig. 5; Table 2). Eg5-CL8 that partitioned with the microtubules at high ionic strength had maximal binding occurring at a tubulin to kinesin molar ratio of approximately 1 (High IS, *s* = 1.2 ± 0.2). At low ionic strength, maximal binding was achieved at a molar ratio suggesting approximately 3 motors bound per tubulin dimer (Low IS, *s* = 3.3 ± 0.9) (Fig. 3a). In addition, under increasing Eg5-CL8 to microtubule ratios above stoichiometric ratios, Eg5-CL8 shows a saturation of binding to the microtubule 7.1 ± 1.5 motors per tubulin dimer (Fig. 4). This was like Cin8-WT, that was previously found to bind to the microtubule with between 3 and 5 Cin8 motors per tubulin dimer (14). With an added MBP tag, Eg5-CL8-MBP showed only apparent stoichiometric binding, with motor domain binding saturating at approximately 1 motor per tubulin dimer (Figs. 4 and 5). We hypothesize that this stoichiometric binding was caused by the large MBP tag inhibiting the super-stoichiometric binding because of steric hindrance (26).

**Figure 5.**
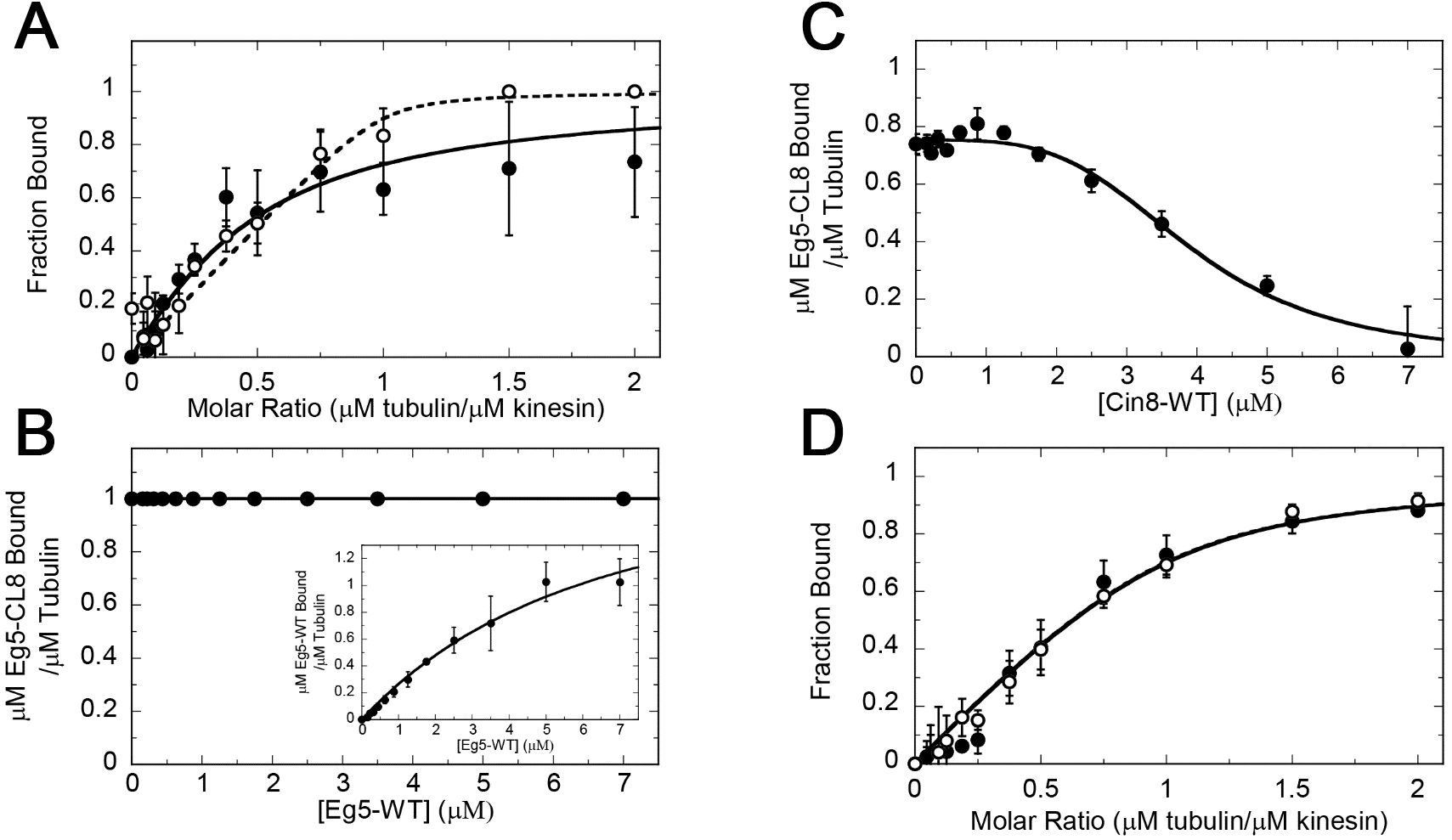
Cosedimentation and competition assays of Eg5-CL8 reveal super-stoichiometric binding and noncanonical binding to the microtubule. (**A**) Cosedimentation of Eg5-CL8 (tagless) at low (closed circles) and high (open circles) ionic strength. Final concentrations are 2 μM Eg5-CL8 (tagless), 0-8 μM MTs, 5 mM ATP. Low IS, *K_d,app_* = 0.8 ± 1.4 μM and s = 3.3 ± 3.5. High IS, *K_d,app_* = 0.04 ± 0.05 μM and s = 1.2 ± 0.2. (B) Plot of average fraction Eg5-CL8 (tagless) bound to the microtubule at increasing Eg5-WT concentrations. *Inset* shows the concentration of Eg5 bound to the microtubule as Eg5 concentration was increased. Final concentrations are 2 μM Cin8-EL8 (tagless), 2 μM MTs, 0-7 μM Eg5-WT, 5 mM ATP. (**C**) Plot of average fraction Eg5-CL8 (tagless) bound to the microtubule at increasing Cin8-WT concentrations. Final concentrations are 1 μM Cin8-EL8 (tagless), 1 μM MTs, 0-7 μM Cin8-WT, 5 mM ATP. (D) Cosedimentation of Eg5-CL8-MBP at low (closed circles) and high (open circles) ionic strength. Final concentrations are 2 μM Eg5-CL8-MBP, 0-4 μM MTs. Low IS, *K_d,app_* = 0.1 ± 0.08 μM. High IS, *K_d,app_* = 0.1 ± 0.02 μM.

Eg5-CL8 displays tight microtubule binding at high ionic strength (*K_d,app_* = 0.04 ± 0.004 μM), and binds approximately 20 times weaker at lower ionic strength (*K_d,app_* = 0.8 ± 0.5 μM) (Fig. 5a; Table 2), which was in contrast to what was expected with a super-stoichiometric *s*-value at low ionic strength and only an apparent stoichiometric *s*-value at high ionic strength. Cosedimentation competition assays between Eg5-CL8 and Eg5-WT were done to check if Eg5-CL8 was capable of binding to a noncanonical site like Cin8-WT (Fig. 5b). Under increasing concentrations of Eg5-WT (up to a 3.5 excess in relation to the microtubule and Eg5-CL8), Eg5 was unable to displace any of the Eg5-CL8 that was in a 1:1 ratio with the microtubule, even when Eg5 was observed to be going onto the microtubule up to the expected saturation of 1 Eg5 motor per tubulin dimer (Fig. 5b). This suggests that Eg5-CL8 was capable of binding to a noncanonical microtubule binding site.

Cosedimentation competition assays between Eg5-CL8 and Cin8-WT were performed to determine whether Eg5-CL8 could compete with Cin8 for the same noncanonical binding site. Increasing concentrations of Cin8 could completely displace Eg5-CL8 from the microtubule when in excess to microtubules and Eg5-CL8 at a ratio of approximately 7:1 (with Eg5-CL8 and MTs in a 1:1 ratio with each other) (Fig. 5c). These results, in combination with Eg5 being unable to displace Eg5-CL8 from the microtubule and an *s*-value that indicated up to 3 motors bound per tubulin dimer, strongly suggests that Eg5-CL8 binds to the same noncanonical binding site as Cin8, and displays the same type of super-stoichiometric binding behavior as Cin8-WT.

### The loop-8 insert from Kip1 does not enable super-stoichiometric binding in Eg5-KL8

The *K_d,app_* for microtubules and the stoichiometry coefficient (*s*) were determined for Eg5-KL8 via cosedimentation assays. Eg5-KL8 displayed tight microtubule binding at high ionic strength (*K_d,app_* = 0.1 ± 0.06 μM), and bound to microtubules 5 times weaker at lower ionic strength (*K_d,app_* = 0.5 ± 0.1 μM) (Table 2). Eg5-KL8 partitioned with microtubules at high ionic strength had maximal binding occurring at a tubulin to kinesin molar ratio of approximately 1 (*s* = 1.3 ± 0.4) (Fig. 6). This was the same stoichiometric behavior that was seen with Cin8-KL8 in our previous paper, and suggests that the loop-8 insert for Kip1 was not sufficient for enabling super-stoichiometric binding.

**Figure 6.**
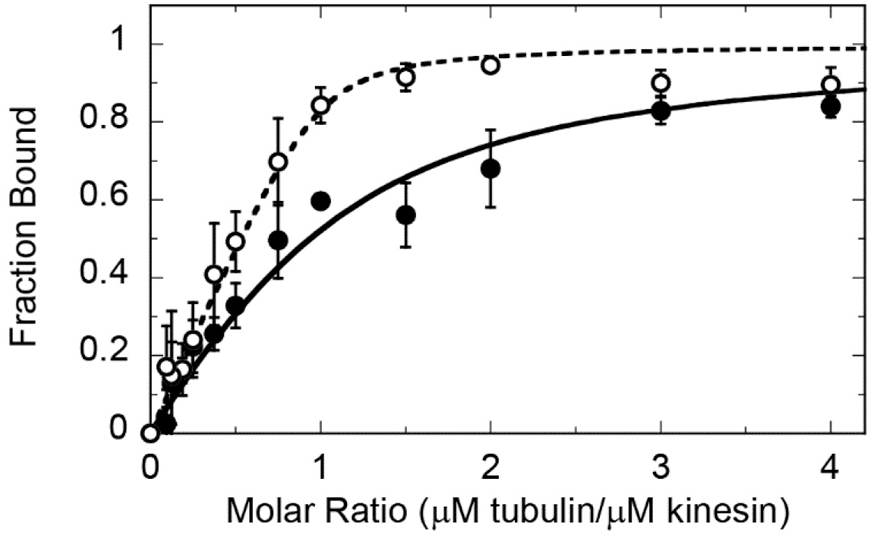
Eg5-KL8 shows Eg5-like stoichiometric binding to the microtubule. Cosedimentation of Eg5-KL8 at low (closed circles) and high (open circles) ionic strength. Final concentrations are 2 μM Eg5-KL8, 0-8 μM MTs. Low IS, *K_d,app_* = 0.44 ± 0.06 μM. High IS, *K_d,app_* = 0.1 ±0.05 μM,s= 1.3 ±0.1.

### The number of Cin8 motors bound per tubulin dimer was affected by ATP

The effect of ATP on super-stoichiometric binding of Cin8 was previously investigated using 1 mM ATP (14), but there was no obvious difference compared to Cin8 binding under no added nucleotide (Cin8-WT, 1 mM ATP: Low IS, s = 3.7 ± 2.1. High IS, s = 6.1 ± 4.8). After observing the effect that high concentrations of ATP (5 mM) had on the solubility of Eg5-CL8 as well as the difference in *k_cat_* for Eg5-CL8 under 5 mM ATP concentrations, we measured microtubule binding via cosedimentation assays of Cin8-WT under 5 mM ATP (Fig. 7). Cin8-WT showed a lower number of Cin8 motors bound per tubulin dimer in the presence of high ATP at low ionic strength (s = 2.2 ± 0.1), than it did without ATP at either low (s = 5.1 ± 4.2) or high (s = 4.0 ± 3.3) ionic strength (Table 2) (14), strongly suggesting that the super-stoichiometric binding behavior of Cin8 was in some way dependent on the presence or absence of ATP.

**Figure 7.**
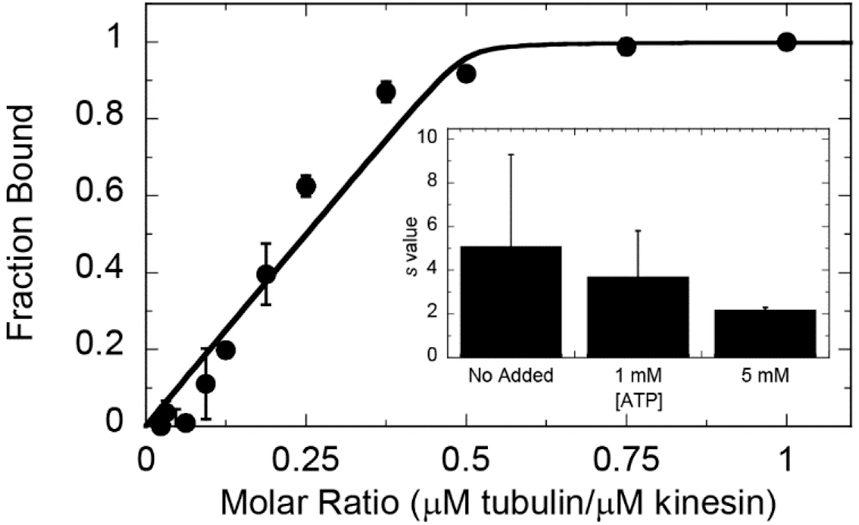
Cosedimentation assays demonstrate fewer Cin8 motors interacting with the microtubule under high ATP conditions. Cosedimentation of Cin8 at low ionic strength with 5 mM ATP. Final concentrations are 2 μM Cin8-trx, and 0-4 μM MTs. *K_d,app_* = 0.007 ± 0.01 μM and *s* = 2.2 ± 0.2. (*Inset*) *s* value for Cin8-WT under 3 different ATP concentrations, no added nucleotide, 1 mM ATP and 5 mM ATP.

Cin8-WT still maintained tight microtubule binding under high ATP conditions (*K_d,app_* = 0.007 ± 0.004 μM), similar to binding observed at low ionic strength without ATP (*K_d, app_* = 0.001 ± 0.003 μM). Even with the increased ionic strength contributed by the ATP (approximately 7.5 mM ionic strength contributed by 5 mM ATP), it did not display any significant weakening of microtubule binding as seen at high ionic strength conditions with no added ATP (*K_d,app_* = 0.1 ± 0.4 μM) (Table 2), which further supports that reduced super-stoichiometric binding was likely caused by the ATP itself, and not a change in ionic strength. This suggests that super-stoichiometric binding to the microtubule was regulated by ATP.

## DISCUSSION

In this study, we utilized molecular biology and biochemical approaches to dissect the mechanism for how the large loop-8 insert in yeast kinesin-5 Cin8 modulates microtubule binding. To this end, we cloned, expressed and purified chimeric motors of Cin8, Eg5 and Kip1 in which their loop-8 inserts were swapped, Cin8-EL8 and Eg5-KL8 showed only canonical binding, while Eg5-CL8 showed binding to both the canonical and noncanonical microtubule binding sites, but with weakened oligomerization.

### Cin8’s loop-8 insert regulates the turnover number and microtubule binding

The two Eg5 mutants (Eg5-CL8 and Eg5-KL8) both have Eg5-like *k_cat_* rates, with minimal difference in *k_cat_* based on ionic strength, suggesting that the Eg5 *k_cat_* was likely not affected by the loop-8 inserts or ionic strength. Cin8-KL8 shows no difference in *k_cat_* compared to Cin8-WT, but Cin8-EL8 has an increased *k_cat_* compared to Cin8-WT (at low ionic strength). This suggests the loop-8 insert from the faster Eg5 motor caused an increased *k_cat_* in Cin8-EL8, while the loop-8 insert from Kip1 does not. These results suggest Cin8 and Kip1 loop-8 inserts may have similar interactions with the Cin8 motor domain that affect their *k_cat_* rates in the same way. Additionally, ionic strength appears to regulate the Cin8-EL8 motor’s *k_cat_*, with its *k_cat_* being faster at low ionic strength conditions, whereas Cin8-WT does not show this affect. This could indicate that Cin8’s loop-8 insert also plays a role in monitoring ionic strength conditions for regulation of the Cin8 motor’s ATPase activity.

Eg5-WT and Cin8-WT both have weakened MT binding at high ionic strength which is expected for kinesin motors that interact with the microtubule through a primarily ionic interaction (14,27–29). Cin8-EL8 showed this same trend, however Eg5-CL8 and Eg5-KL8 both showed tighter microtubule binding at high ionic strength instead of weaker binding. This could suggest that both the Cin8 and Kip1 loop-8 inserts have similar effects on microtubule binding. An increase in hydrophobic interactions between the motor domain and the microtubule could explain why higher salt concentrations, which can drive hydrophobic residues to interact more tightly (30–32), may cause increased microtubule binding, rather than weakening it like is normally seen in the traditional, kinesin-microtubule interaction. For this to occur with only the Eg5 mutants and not the wild type Cin8 motor, suggests that at the Cin8 loop-8 insert (and possibly the Kip1 loop-8 insert) has interactions with its wild type core motor domain that affects its microtubule binding interactions and which are likely not preserved when placed onto the Eg5 motor domain. Cin8’s loop-8 insert has many residues, and is predicted to be disordered, so determining which residues within the loop-8 may be causing this effect on microtubule binding is difficult. However, Kip1’s loop-8 insert is the same length as Eg5’s loop-8 insert, at only 7 amino acids, and moreover it has a very different sequence of amino acids (NNNNNSS) when compared to Eg5 (DPRNKRG), with the residues in Kip1’s loop-8 insert being polar, but not charged like in Eg5’s loop-8. These less charged residues may cause the increased hydrophobic effect seen with the Eg5-KL8 motor. Additionally, Cin8-EL8 shares a similar 7NS region in its loop-8 (NNSSSNS) that could be interacting with the microtubule similarly to the Kip1 loop-8.

### Loop-8 plays a significant role in the novel super-stoichiometric binding of Cin8, and in regulating the traditional stoichiometric binding of Eg5

We utilized a combination of classic cosedimentation and competition cosedimentation assays with two mutant motor domain constructs, Cin8-EL8 and Eg5-CL8, to investigate the role of loop-8 in super-stoichiometric binding observed in the wild type monomeric Cin8 motor domain (14). The well-studied Eg5-WT and Cin8-WT motor domains were used as controls for typical canonical and noncanonical microtubule binding, respectively. The mutant constructs showed microtubule binding stoichiometry coefficients (*s*) like those of the wild type motor that provided their loop-8 inserts. Cin8-EL8 shows only stoichiometric and traditional canonical microtubule binding. Similarly, Eg5-CL8 shows super-stoichiometric binding, as well as an ability to bind to the same noncanonical binding site as the wild type Cin8 motor. This strongly suggests that the loop-8 insert was necessary for the novel noncanonical binding of Cin8 that was previously characterized (14).

### Loop-8 required for noncanonical binding, but not dimerization

While Cin8’s loop-8 insert was required for ~4:1 binding of Cin8 to the microtubule, and enables noncanonical binding of Eg5-CL8, it may not be the only requirement for all super-stoichiometric binding. We have postulated previously that there are two mechanisms by which Cin8’s super-stoichiometric binding occurs, one being the binding to a noncanonical site, and the other being dimerization of the motor domains to each other on either the canonical or noncanonical sites (14). For both mechanisms, up to four motor domains could bind per tubulin dimer in the microtubule lattice. The competition assays between the mutant and the wild type Cin8 and Eg5 motor domains indicated that Cin8’s loop-8 insert was responsible for noncanonical binding since any construct lacking the large loop-8 insert can be competed off by canonical binding Eg5, whereas motors that possess Cin8’s loop-8 insert can only be fully competed off by Cin8-WT. However, these competitions also showed a strong sigmoidal character to them, suggesting that there may be still some type of dimerization occurring between the motor domains. Dimerization of Eg5-CL8 was not unexpected since this construct showed super-stoichiometric binding in the other experiments. The sigmoidal nature of the Cin8-EL8 construct was unexpected though. Cin8-EL8 appears to bind only stoichiometrically in other experiments, however in the competition assays with Eg5-SCRK, the sigmoidal curve suggests that some of the Cin8-EL8 motors may still interact with each other in a way that leaves empty sites for the Eg5-SCRK to bind to without displacing the Cin8-EL8. This may be an indication of some dimerization between the Cin8-EL8 motors, even without the loop-8 insert and could suggest that the Cin8 motor cores can interact with each other, or were interacting via regions of the loop-8 outside the loop-8 insert region.

### Loop-8 insert from Kip1 was insufficient for super-stoichiometric binding and super-stoichiometric binding was not a requirement for bidirectionality

Both Kip1 and Cin8 are bidirectional, but only Cin8’s loop-8 insert appears to enable super-stoichiometric binding (9–12). The Cin8-KL8 construct characterized in our earlier studies (previously referred to as Cin8-Δ99 (10,14)), as well as the Eg5-KL8 studied in this study showed only stoichiometric binding to the microtubule, suggesting that Kip1’s loop-8 insert was insufficient for super-stoichiometric binding, despite Kip1 being another bidirectional motor. Furthermore, Cin8-KL8 was shown to still possess bidirectional motility (10). The fact that the bidirectional Cin8-KL8 does not bind super-stoichiometrically indicates that super-stoichiometric binding is not an absolute requirement for bidirectionality.

### Loop-8 may contain a secondary ATP binding site that regulated super-stoichiometric binding

The monomeric Eg5 motor domain was highly soluble in solution and can be concentrated to above 500 μM (25). However, Eg5-CL8 precipitated out of solution under high concentrations (40 μM), yet could be re-solubilized with the addition of high concentrations ATP (5 mM). This indicated that the addition of Cin8’s loop-8 insert to the normally soluble Eg5 motor domain not only caused the motor domain to precipitate, but that the presence of ATP can re-solubilize it, strongly suggesting that the ATP has a direct interaction with the Cin8’s loop-8 insert. Additionally, at 1mM ATP the ATP concentration was well above the *K_m,ATP_* for Cin8 or Eg5. At concentrations above 1 mM, a large majority of the active sites would already be saturated with ATP, but at 5 mM ATP we did observe a significant effect on the binding stoichiometry of Cin8-WT with microtubules. With no added nucleotide. Cin8 binds around 4:1 motors per tubulin dimer, however at 5 mM, the number of motors per tubulin dimer has dropped to 2:1. This strongly suggests that there is a second ATP binding site with a much lower binding affinity, and that this site has a regulatory effect on super-stoichiometric binding.

### Potential in vivo function of the kinesin-5 loop-8 insert

The loop-8 insert for Cin8 was required for noncanonical binding, but super-stoichiometric binding was not required for bidirectionality. However, super-stoichiometric binding may still play a role in regulating bidirectional motility in the cell. We have hypothesized previously that Cin8’s super-stoichiometric binding occurs via both noncanonical binding and dimerization of the motor domains (14). There have also been two different mechanisms found to affect bidirectionality in kinesin-5 motors: a clustering model (33) and a crowding model (9,26). We hypothesize that the two different mechanisms of super-stoichiometric binding are involved in different mechanisms of regulating bidirectionality. Cin8’s noncanonical binding could provide a crowing mechanism by bringing the motor heads into high local concentration to induce directional switching. Additionally, dimerization of Cin8 motors could enable the formation of Cin8 clusters at the minus-ends of microtubules (33). Further studies well be needed to confirm whether these physiological roles of Cin8’s super-stoichiometric binding are correct

## EXPERIMENTAL PROCEDURES

### Experimental conditions

Experiments were conducted at 298 K in ATPase buffer (20 mM HEPES, pH 7.2 using sodium hydroxide, 2 mM magnesium chloride, 20% (v/v) glycerol) unless otherwise noted. Typically, potassium chloride at 7.5 mM (referred to as low ionic strength) or 75 mM (referred to as high ionic strength) was added to the ATPase buffer. For ionic strength determination, the buffer contributed 7 mM and magnesium chloride contributed 6 mM plus the molarity of the monovalent potassium chloride added to the ATPase buffer. For biochemistry experiments using microtubules, 20 μM paclitaxel was added to the ATPase buffer to stabilize microtubule polymers.

### Cloning

The oligo nucleotides used for cloning each construct are listed in Table 3. For cloning the Eg5-CL8-MBP (MW = 95.8 kDa) construct, the sequence corresponding to the 99-residue loop-8 insert of Cin8 was amplified via polymerase chain reaction (PCR) using oligos that contained homologous sequence to the Cin8 loop-8 insert and flanking homologous sequence of the Eg5 motor domain (ECL8-1 and ECL8-4). The two halves of the Eg5 gene on either side of loop-8 insert were amplified with oligos that contained overlap regions matching the Cin8 loop-8 region (EG5-1 and ECL8-2 for the 5’-end of the gene and EG5-2 and ECL8-3 for the 3’-end). The amplified Cin8 loop-8 insert and the Eg5 motor domain segments were fused together using several rounds of double fusion PCR. The completed Eg5-CL8 sequence was inserted into a modified pET16b vector, and a maltose binding protein tag was amplified from a pMALc5x vector and inserted into the Eg5-CL8/pET16b vector (construct from N-terminus is MBP-His_6_-TEV-Eg5-CL8).

**TABLE 3.**
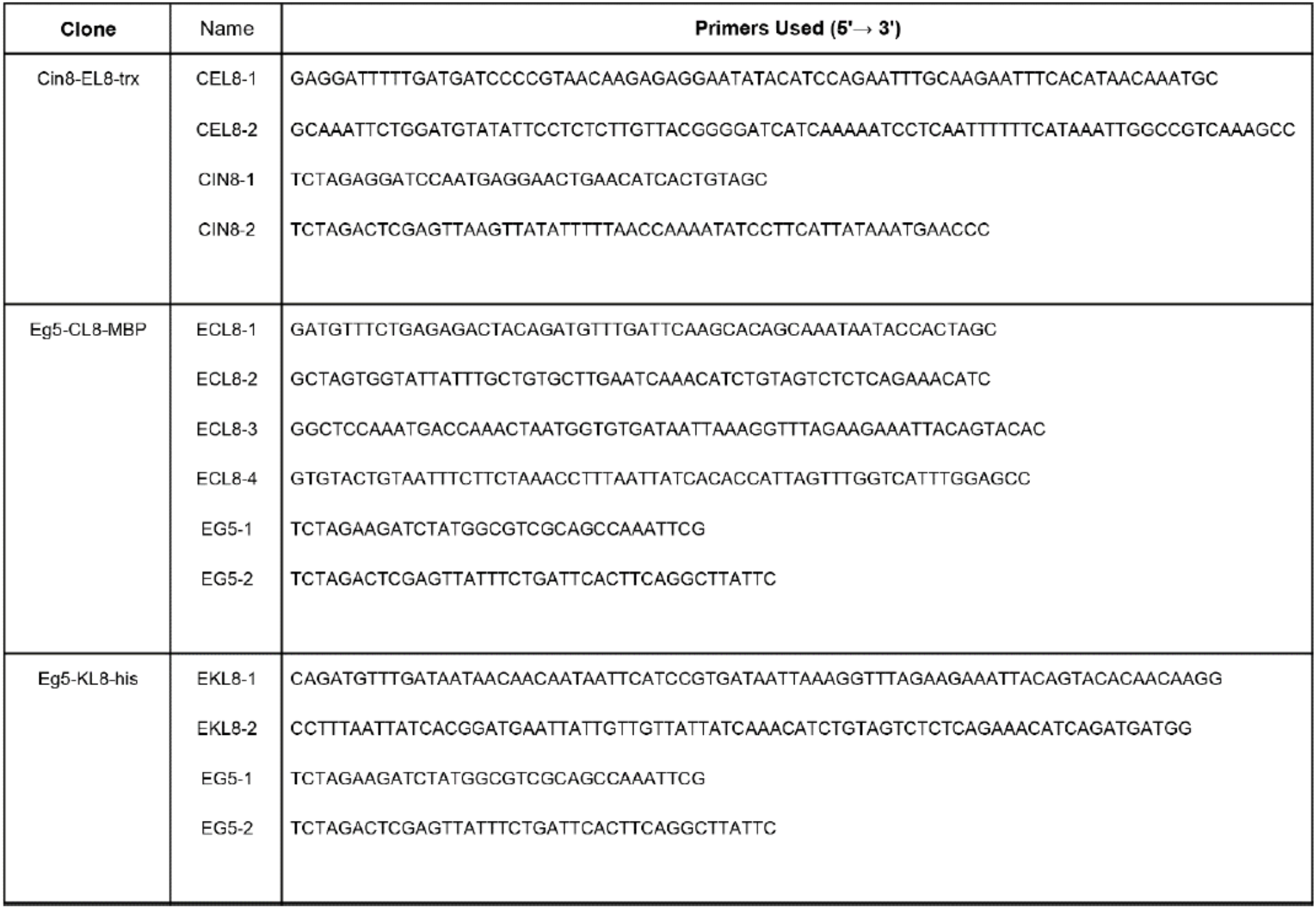
Oligo nucleotides used for cloning chimeric kinesin-5 constructs.

For the Cin8-EL8-trx (MW = 56.0 kDa) and Eg5-KL8-his (MW = 43.3 kDa) constructs, overlapping primers containing the loop-8 sequences for Eg5 or Kip1 flanked by sequences matching the flanking Cin8 motor domain sequences (Eg5 loop-8 insert were CEL8-1 and CEL8-2; Kip1 loop-8 insert were EKL8-1 and EKL8-2) were used along with oligos matching the 5’-end (CIN8-1 and EG5-1) and 3’-end (CIN8-2 and EG5-2) of the motor domain sequence for Cin8 or Eg5, respectively. The fusion motor genes were amplified in two halves around the inserted loop-8 sequence, and then those halves were combined using double fusion PCR to insert the 7 amino acid sequences into the constructs. The Cin8-EL8 sequence was then transferred into a modified pET32b vector (pJCC04a) for bacterial expression to achieve an N-terminal thioredoxin tag and TEV cleavage site (34). The Eg5-KL8 sequence was transferred into a modified pET16b vector for bacterial expression to achieve an N-terminal His_6_ tag and TEV cleavage sequence.

### Protein expression and purification

Wild type Cin8 (Cin8-WT) motor constructs (Cin8-his and Cin8-trx) were expressed and purified as previously described (14). Each chimeric construct (Eg5-CL8-MBP, Cin8-EL8-trx and Eg5-KL8) was purified by a similar method, but were purified using nickel-nitrilotriacetic acid resin (Qiagen, Germantown, MD). The final protein concentration was 15-260 μM (~2-10 mol ADP per mol kinesin). Expression and purification of nucleotide-free human kinesin-5 (Eg5-WT) was performed as previously described (22,23,25,35). Microtubules were prepared using an aliquot of purified bovine brain tubulin that was thawed and cycled, and the microtubules were stabilized with 50 μM paclitaxel (Thermo Fisher Scientific, Waltham, MA) as previously described (14). Each kinesin preparation was 90-95% pure via densitometry analysis of Coomassie stained SDS gels, except Eg5-CL8-MBP and Eg5-CL8 (tagless) which both showed additional bands consistent with partial expression of the MBP tag. We removed purification tags from the Eg5-CL8-MBP and Cin8-EL8-trx constructs using TEV protease to create ‘tagless’ Eg5-CL8 (MW = 50.7 kDa) and Cin8-EL8 (MW = 41.5 kDa) for testing the untagged motor domain behavior in comparison to the tagged variants. Protein concentration was determined using via Bradford assay reagent (Thermo Fisher Scientific, Waltham, MA) with bovine serum albumin (BSA) as the standard, and by densitometry analysis of Coomassie stained SDS gels.

### Steady-state ATPase assay

The basal and microtubule-stimulated ATPase activities of kinesin-5 motors were measured by the NADH coupled assay as previously described (14,27). For ATP-dependent experiments, the initial ATPase velocity was plotted against the ATP concentration and the data were fit to the Michaelis-Menten equation 1, where [E_0_] is the total concentration of kinesin, *k_cat_* is the maximum rate of ATP turnover per kinesin active site at infinite [ATP], and *K_m,ATP_*, is the Michaelis constant, which is defined as the [ATP] that yields ½[E_0_]*k_cat_*. For microtubule-dependent experiments, the initial ATPase velocity was plotted against the microtubule concentration, and the data were fit to the quadratic equation 2, where [E_0_] is the total kinesin motor concentration, *k_cat_* is the maximum rate of ATP turnover per kinesin active site at infinite microtubule concentration, *K*_0.5,MT_ is the microtubule concentration that yields ½[E_0_]*k_cat_*, and *k*_basal_ is the experimentally determined basal rate of ATP turnover at zero microtubule concentration. Each plot corresponds to the average ATPase rate normalized to 1 μM motor for n = 3-5 experiments, with error bars showing the standard deviation of each data point when the experiments were replicated. Unless otherwise stated, the reported *k_cat_* values are the values obtained from MT-dependent steady-state experiments.

### Cosedimentation assays

Purified kinesin with no added nucleotide was incubated with paclitaxel-stabilized microtubules for 15-60 min and microtubule·kinesin complexes were pelleted by centrifugation in a Beckman Optima L-100 XP ultracentrifuge using the Ti-42.2 fixed-angle rotor at 100,000xg for 30 min at 298 K as described (14,27). For Cin8-trx, Cin8-EL8 (tagless), Eg5-CL8 (tagless), Eg5-CL8-MBP and Eg5-KL8, gel samples were prepared for the supernatant and pellet fractions at equal volumes, and the proteins were resolved by SDS-PAGE and stained with Coomassie brilliant blue R-250. The fraction of total kinesin bound to microtubules was plotted against the microtubule concentration, and the data were fit to quadratic equation 3, where [E_0_] is the total kinesin concentration, *K_d,app_* is the apparent *K*_d,MT_ that represents the average binding constant for all binding sites, and *s* was the stoichiometry coefficient to quantify the number of binding sites. Error bars correspond to the standard deviation obtained from n = 2-6 experiments.

### Competitive cosedimentation assays

A series of competition experiments were designed to characterize the microtubule binding behavior of Cin8-EL8 and Eg5-CL8 by displacing them from the microtubule with canonical binding kinesins Eg5-WT or Eg5-SCRK (an ATPase dead Eg5 mutant, (25)) and noncanonical binding kinesin Cin8-trx (14). The mutant kinesin (designated kinesin A in equation 4) and competing kinesin (kinesin B in equation 4) were incubated with microtubules in the same reaction. The mixture was equilibrated at 298 K for 60 min before centrifugation. Data were fit to equation 4 (36), where is the fraction of mutant kinesin A bound to the microtubules, [kinesin B] is the concentration of the competing kinesin B, y_t_ is the maximal concentration of kinesin A bound per tubulin dimer, K_i_ is the concentration of kinesin B needed to compete off half of the total kinesin A from the microtubule, and *m* is cooperativity coefficient. Error bars correspond to the standard deviation obtained from n = 3 experiments. For competition experiments between Cin8-EL8 and ATPase-deficient Eg5-SCRK, which both have molecular weights that cannot be resolved from tubulin on an SDS-PAGE gel, the concentrations of Cin8-EL8 in the supernatant was measured using the NADH coupled assay. A supernatant sample was mixed with an equal volume of 2x NADH reaction mix (0.5 mM phosphoenolpyruvate, 0.4 mM NADH, 5 U/mL rabbit pyruvate kinase, 8 U/mL lactic dehydrogenase, 1 mM ATP, 5 μM tubulin as microtubules). The fraction of Cin8-EL8 bound (*f_b_*) at each condition was determined by equation 5.

## EQUATIONS

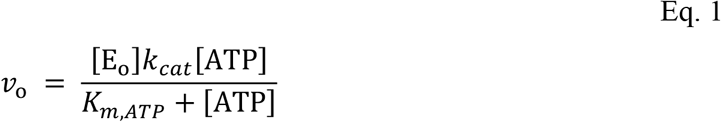

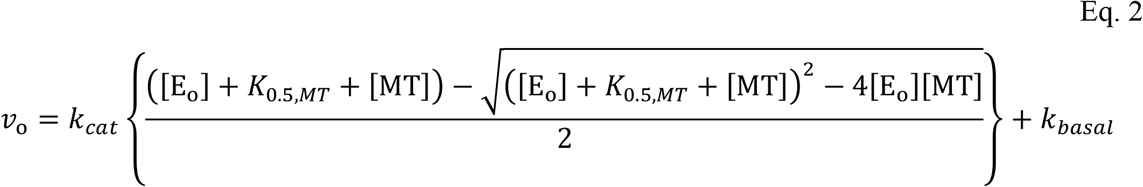

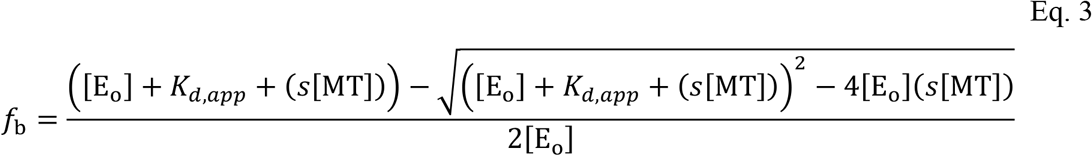

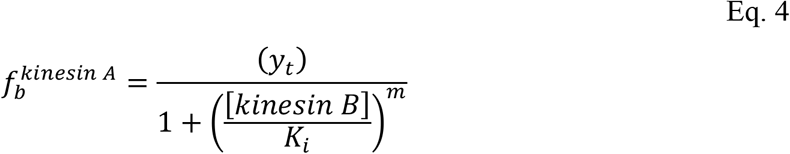

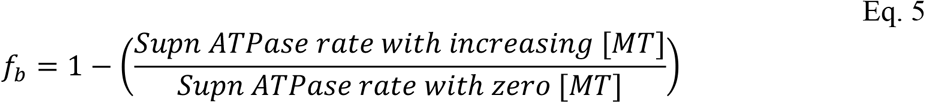

## ACKNOWLEDGEMENTS

We thank Benjamin Walker for valuable discussions. This work was supported by Indiana University and the National Science Foundation (MCB 1614514 to J.C.C.).

## AUTHOR CONTRIBUTIONS

K.M.B. and J.C.C. designed the study, performed the research, and wrote the manuscript.

